# Differential effects of open- and closed-loop intracortical microstimulation on firing patterns of neurons in distant cortical areas

**DOI:** 10.1101/534032

**Authors:** Alberto Averna, Valentina Pasquale, Maxwell Murphy, Maria Piera Rogantin, Gustaf Van Acker, Randolph J. Nudo, Michela Chiappalone, David Guggenmos

## Abstract

**Background:** Intracortical microstimulation can be used successfully to modulate neuronal activity. Activity-dependent stimulation (ADS), in which action potentials recorded extracellularly from a single neuron are used to trigger stimulation at another cortical location (closed-loop), is an effective treatment for behavioral recovery after brain lesion in rodents. Neurophysiological changes in cortical communication induced by ADS, and how these changes differ from those induced by open-loop random stimulation (RS) are still not clear.

**Objectives:** We investigated the ability of ADS and RS to induce changes in firing patterns in distant populations of neurons in healthy anesthetized rats.

**Methods:** For this study we used 23 adult Long-Evan rats, recording from a total of 591 neuronal units. Stimulation was delivered to either forelimb or barrel field somatosensory cortex, using either randomly-timed stimulus pulses or ADS triggered from neuronal spikes recorded in the rostral forelimb area (RFA) of the motor cortex.

**Results:** Both RS and ADS stimulation protocols rapidly altered spike firing within RFA compared with no stimulation. Changes consisted of increases in mean firing rates and patterns of spike firing as measured by the revised Local Variation metric. ADS was more effective than RS in increasing short-latency evoked spikes during the stimulation periods, by producing a reliable, progressive increase in stimulus-related activity over time.

**Conclusions:** These results are critical for understanding the efficacy of electrical microstimulation protocols in altering activity patterns in interconnected brain networks. These data further strengthen the idea that activity-dependent microstimulation, can be used to modulate cortical state and functional connectivity.

## Introduction

Focal, invasive electrical stimulation techniques such as intracortical microstimulation (ICMS) and deep brain stimulation (DBS) have the ability to target and directly activate a relatively small population of neurons compared to non-invasive techniques such as transcranial magnetic stimulation (TMS) or transcranial direct current stimulation (tDCS). While ICMS is typically employed in invasive animal studies, focal properties of invasive electrical stimulation have made DBS advantageous for implementation into clinical tools for treating a variety of neurological conditions such as epileptic seizures [1–4] and Parkinson’s disease [5–9]. ICMS, specifically, is also being investigated for the direct treatment of conditions such as pain and depression among others, and in brain-computer interfaces to augment or restore lost function after injury, such as for visual [10–14] or somatosensory [15–17] neuroprosthetic devices. While the local physical and neurophysiological effects of this type of stimulation have been described in detail [18–22], there is much less information on how focal stimulation impacts areas distant from the immediate spread of the electrical current or across multi-synaptic pathways. This is an important consideration, as these stimulated regions are not isolated [23] but have anatomical connections with several other regions within the brain. The effectiveness of treatments that rely on focal electrical stimulation are likely dependent on the modulation of these pathways but there are still open questions about how this stimulation affects the firing patterns within these distant regions.

The impact of focal electrical stimulation on distant brain regions may be even more pertinent in “closed-loop” designs, in which intrinsic neural activity drives stimulation protocols. One such design, named activity-dependent stimulation (ADS), uses the occurrence of action potentials (spikes) in one neuron to trigger stimulation at another location or electrode site. ADS relies on the concept of Hebbian plasticity, in which repeated concomitant firing of two neurons will strengthen the connection between them. In closed-loop systems, secondary neuronal firing is induced via the stimulation. By artificially pairing spike-firing in one populations of neurons with focal electrical stimulation of a second population of neurons, it may be possible to shape the efficacy of specific neural pathways *in vivo* [24–28].

The purpose of the present study was to determine if neural firing patterns in distant regions are altered in response to short-duration stimulation sessions in anesthetized rats. To model a relevant system *in vivo*, the rostral forelimb area (RFA), a premotor area, was used for neural recordings while either closed-loop ADS or random stimulation (RS) was delivered to somatosensory cortex (S1), either in the S1 forelimb area (S1FL) or the S1 barrel field (S1BF). These areas share reciprocal neuroanatomical connections, providing an anatomical framework for changing synaptic efficacy [29]. In a previous study, we found that an ADS protocol in a chronic injury model, i.e., pairing the occurrence of spikes in RFA with ICMS applied to S1, led to increased firing within RFA over a period of several days [28]. In the present study, we investigated ADS effects under the more controlled conditions of an anesthetized preparation, enabling us to assess alterations in RFA spike firing patterns over single stimulation sessions. This preparation allowed us to more readily test various parameters such stimulation condition (ADS or RS) and stimulation location (S1FL or S1BF). Because these regions are functionally and anatomically connected, the results may be generalizable to other, similarly interconnected regions. Understanding changes in neuronal activity in distant areas in response to focal closed-loop and random ICMS may lead to more effective brain stimulation protocols that target functional connectivity within specific brain pathways, and thus, to improve current therapeutic devices for neurological disorders.

## Methods

### Animals

All experiments were approved by the University of Kansas Medical Center Institutional Animal Care and Use Committee. A total of 23 adult, male Long-Evans rats (weight: 350-400 g, age: 4-5 months; Charles River Laboratories, Wilmington, MA, USA) were used in this study.

### Surgical Procedures

Prior to surgery, anesthesia was induced with gaseous isoflurane within a sealed vaporizer chamber followed by bolus injections of ketamine (80-100 mg/kg IP) and xylazine (5-10 mg/kg). Anesthesia was maintained throughout the procedure with repeated bolus injections of ketamine (10-100 mg/kg/hr IP or IM) as needed. A midline incision was made to expose the skull. A laminectomy was performed at the cisterna magna to drain cerebrospinal fluid, thus controlling brain edema. A craniectomy was made over the extent of the location of pre-motor cortex (PM or RFA), primary motor cortex (M1 or CFA) and primary somatosensory cortex (S1) using a drill with a burr bit. Saline was applied periodically to avoid heat generation by the drill. The dura was resected over the extent of the opening. Silicone oil (dimethylpolysiloxane) was applied to the cortical surface to avoid desiccation and facilitate electrophysiological procedures. Following the experiment, rats were humanely euthanized using pentobarbital (390 mg IP).

### Identification of Cortical Areas of Interest

Upon completion of the surgical procedure, a picture of the vascular pattern of the cortical surface was taken and uploaded to a graphics program (Canvas GFX, Inc., Plantation, FL, USA) where a 250 μm virtual grid was overlaid onto the image. The location of RFA was determined using standard ICMS protocols [30]. Briefly, a glass microelectrode (10-25 μm diameter) filled with saline was inserted into the cortex at a depth of ~1700 μm (impedance at 1,000 Hz = ~ 500kΩ). Stimulus trains of 13-200μs pulses at 333 Hz were applied at 1 Hz intervals using a stimulus isolator (BAK Electronics, Umatilla FL, USA). Current was increased until a visible movement about a joint was observed, up to a maximum current of 80μA. Sites were sampled at a resolution of 250μm between locations. RFA was defined as the region in the frontal cortex where ICMS evoked forelimb movements. RFA was bordered caudally by a region where ICMS evoked neck and trunk movements, and medially by face and jaw movements [31]. RFA was easily distinguished from the caudal forelimb area (CFA) based on its relatively smaller size.

Somatosensory areas were identified by correlating neural responses evoked by slight indentation of the skin or vibrissae brushing. A single-shank, sixteen-channel Michigan style electrode (A1×16-5mm-100-703-A16, NeuroNexus, Ann Arbor, MI, USA) was lowered to a maximum depth of ~1700 μm and attached to a unity gain headstage connected to a digitizing pre-amplifier and piped to a processing unit (Tucker-Davis Technologies, Alachua, FL, USA). Activity of all 16 channels was displayed in real time on a computer screen for visual discrimination of spikes, and a user selected channel was sent to a speaker for auditory discrimination of spikes. S1FL was defined as the region where a consistent, short-latency spike discharge was evoked in response to light stimulation of a small receptive field on the wrist, paw or digits. S1BF was defined by a constant short-latency spike discharge evoked by deflection a single or small number of the mystacial vibrissae. For each sensory area, multiple cortical sites were characterized to ensure reliability of the target location.

### Experimental protocol

Following identification of RFA and S1 FL/BF, a four-shank, sixteen-contact site microelectrode probe (A4×4-5mm-100-125-703-A16, NeuroNexus, Ann Arbor, MI, USA) was placed within RFA at a depth of ~1600 μm for recording purposes (Fig. 1A). An active unity gain connector was attached to the probe and connected through a preamplifier for recording (Tucker-Davis Technologies - TDT, Alachua, FL, USA). Passband filtered data (300-5000Hz) underwent online spike detection using a principal component sorting algorithm through which neural data could be sorted from non-cellular data online. An activated single-shank, sixteen-contact microelectrode probe with an impedance of 100-300kΩ (A1×16-5mm-100-703-A16, NeuroNexus, Ann Arbor, MI, USA), was inserted into the somatosensory area to a depth of ~1500 μm for stimulation purposes (Fig. 1A). For all experimental conditions, a single 60 μA balanced biphasic, cathodal-leading stimulation pulse (200 μs positive, 200 μs negative) was delivered into S1 through a single contact site on the electrode (site 6, corresponding to the tip of the electrode) on each stimulation trigger. The decision to limit stimulation amplitude to 60 μA (12nC/phase and charge density of 1.7 mC/cm^2^) was motivated by the finding that stimulation <= 60 μA has been shown to have a negligible effect on the electrode-tissue interface [32] and on the neuronal tissue itself [33]. Stimulation pulses were delivered using a stimulus isolator and a passive headstage (MS16 Stimulus Isolator, Tucker-Davis Technologies - TDT, Alachua, FL, USA).

**Fig. 1.**
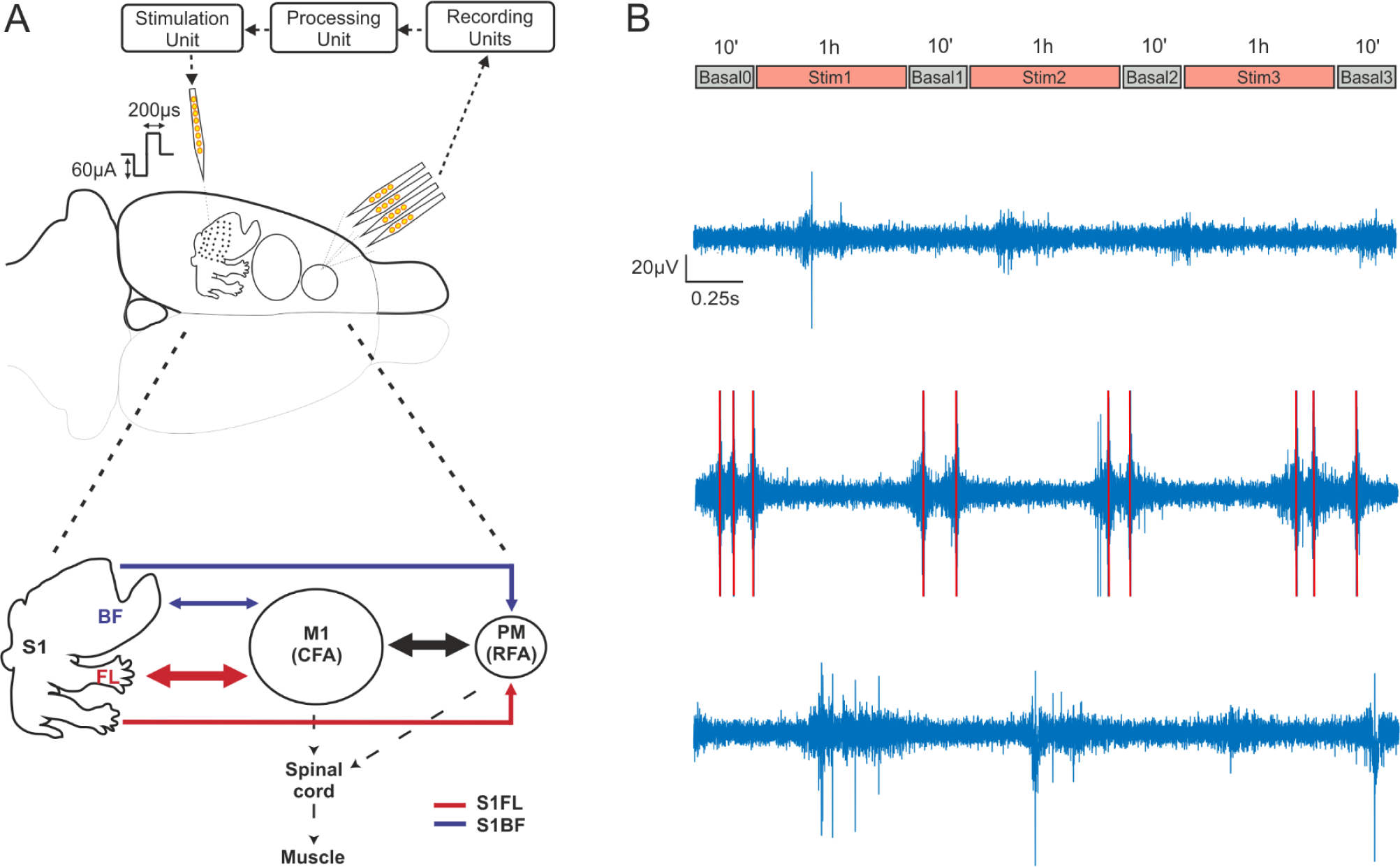
Recording and stimulation experimental paradigm. (A) Extracellular recordings were obtained through a four-shank, 16-contact microelectrode probe in anesthetized rats. Signals were acquired *(Recording Unit)* and processed to detect single-unit activity *(Processing Unit)* selected by the user and employed as reference neuron to trigger a stimulus pulse to a channel on the single-shank microelectrode *(Stimulation Unit*). A summary diagram showing the main cortico-cortical anatomical connections between regions is shown in Fig. 1A, bottom. Arrow thickness corresponds to the amount of labeling (medium or high) suggested by Zakiewicz [53]. (B) Recording sessions (in RFA) consisted of three one-hour intermittent periods of stimulation with either ADS or RS to either S1BF or S1FL, each separated by ten-minute periods of no stimulation. Example traces in a representative experiment of extracellular recording during the first basal period (Basal0) (C), during the first stimulation session (Stim1, D) and during the second basal period, after the stimulation (Basal1, D). Red lines represent the electrical stimulation artifacts.

Rats were randomly assigned to one of four experimental groups based on stimulation condition (activity-dependent stimulation, ADS or random stimulation, RS) and stimulation location (S1 forelimb area, FL or S1 barrel field, BF). Thus, each rat was subjected to only one set of experimental parameters. An additional control group (CTRL) received no stimulation (see Table 1 for group assignments). For all groups, the experimental protocol consisted of four 10-minute periods where the stimulator was set to deliver 0 μA (i.e. Basal0, Basal1, Basal2 and Basal3), interspersed with three one-hour periods where stimulation was delivered at 60 μA (Stim1, Stim 2, Stim3), for a total of approximately three hours and forty minutes of recorded data for a single experimental session (Fig. 1B). Based on group assignment, rats received either random stimulation (RS) pulses (Poisson distribution, mean stimulation interval of 7Hz) or activity-dependent stimulation (ADS, mean stimulation interval of 7Hz), which was triggered on a single spike profile of a channel in RFA with a moderate rate of firing (4-10Hz). For ADS there was a 10ms delay between spike detection and stimulation. There was also an 18ms “blanking” period immediately following the stimulation pulse where stimulation could not be triggered to eliminate any direct stimulation feedback loop.

**Table 1:** Total number of animals and number of units examined per experimental condition.

| Location of the stim |  | Stimulation Type |  |  | Total |
| --- | --- | --- | --- | --- | --- |
| electrode |  | RS | ADS | CTRL |  |
| S1 BF |  | 4 / 113 | 6 / 180 | - | 10 / 293 |
| S1 FL |  | 4 / 103 | 5 / 114 | 4 / 81 | 13 / 298 |
| Total |  | 8 / 216 | 11 / 294 | 4 / 81 | 23 / 591 |

In four rats (1 ADSBF, 2 ADSFL, 1 RSBF) from the total of 23, recording sessions were truncated after the first two hour-long stimulation periods (after Basal2), due to either a substantial decrease in neural activity, presumably due to the anesthetic protocol, or due to technical issues related to environmental noise, but unrelated to the stimulation protocol. However, data from these rats up to the time that the protocol was terminated were considered to be valid and have been included in the analyses. Control experiments underwent the same protocol as ADS, but the stimulator was set at 0 μA for the entire recording session.

### Data Processing

Bandpass-filtered neural data (~300 Hz to 3 kHz) were recorded (Tucker-Davis Technologies - TDT, Alchulta, FL, USA) at ~25 kHz per channel and processed using custom MATLAB (The Mathworks, Natick, MA, USA) scripts. A custom offline spike detection algorithm, Precise Timing Spike Detection [34], was used to discriminate spikes [34] followed by superparamagnetic clustering [35, 36] to sort the detected spikes. A supervised discrimination method was used to singularly validate the spike profiles.

### Mean Firing Rate (MFR)

The neuronal firing rates were evaluated before and after the stimulation by calculating the mean firing rate in each period. Neurons whose firing rate was less than 0.01 spikes/s were discarded. Determination of whether the difference in firing rates between two time-points for a given unit significantly deviated from a null (zero centered) distribution was calculated using a bootstrapping method [37]. The two time segments to be compared (Basal_i_, Basal_i+1_; i=1:3) were divided into 1 min bins, and then randomly shuffled 10,000 times into two groups. The differences between the means of the two randomly shuffled groups produced a null-distribution. The real difference was significant if it fell outside of the 95% confidence interval of the null-distribution.

### Local Variation compensate for Refractoriness

The temporal patterns of spike activity exhibited in RFA was evaluated using a revised version of Lv parameter, namely LvR (Local Variation compensate for Refractoriness), as proposed in [38]. LvR measures the local variation of the ISI and describes the intrinsic firing irregularity of individual neurons, not being confounded by firing rate fluctuations [38]. This metric produces a value, ranging from 0 to more than 2 and can be used to classify the individual neuron’s activity into *Regular* (approx. 0.5±0.25), *Random* (approx. 1±0.25) and *Bursty* (approx. 1.5±0.25) firing patterns [38].

### Post-Stimulus Time Histogram

Post-Stimulus Time Histograms – PSTH [39, 40] (1 ms bins, normalized over the total number of stimulation pulses) of stimulus-associated action potentials of each sorted unit were calculated during the 28ms following stimulus pulses delivered from either S1FL or S1BF. The area under the normalized PSTH curve was used to quantify the total amount of stimulation-evoked neural activity during each stimulation phase.

### Statistical Model

Statistical analysis was performed by applying a general linear mixed effects model for repeated measures analysis and SAS/STAT^®^ software (Version 9.2 of the SAS System for Windows, SAS Institute Inc., Cary, NC, USA). This model was utilized as it allows more flexibility than traditional multivariate regression analysis. It permits modeling of not only the means of the data (as in the standard linear model) but also their variances as well as within-subject covariances (i.e., the model allows subjects with missing outcomes -unbalanced data-to be included in the analysis).

In this model, the subjects were the recorded neurons from each rat *(neuron (rat))*. The model included two fixed between-subject factors: the stimulation condition (with three levels for factor *StimCond:* RS, ADS and CRTL) and the stimulation location or *Area* (with two levels for factor *Area*: S1FL and S1BF). The outcome was a variable measured at four fixed time points (within-subject factor *Time*). The model also included the interactions of second and third order among the factors (*StimCond * Area, StimCond * Time, Area * Time* and *StimCond * Area * Time*).

The rat was considered a random factor and implied a different intercept for each rat. The variance-covariance matrix for errors was considered unstructured, as it was different for each subject. The MIXED procedure of SAS, by default, fits the structure of the covariance matrix by using the method of restricted maximum likelihood (REML) [41], also known as residual maximum likelihood. Finally, the fixed part of the expected values of the basal period recorded at the time *t*, with the stimulus *s* applied to the area *a*, was: μ + τ_t_ + α_*a*_ + σ_*s*_ + δ_*t,a*_ + γ_*t,s*_ + ε_*a,s*_ + γ_*t,a,s*_, where the parameters τ_*t*_, with *t ∈ {1,2,3,4}*, referred to factor *Time*, the parameters α_a_,with *a ∈ {FL, BF}*, referred to factor *Area*, the parameters σ_s_,with s ∈ {*ADS, RS, CTRL*}, referred to factor *StimCond* and the parameter δ_*t,a*_, γ_*t,s*_, ε_*a,s*_ and γ_*t,a,s*_ referred to second and third order interactions of. The reference level of the factor *StimCond* was set to be CTRL first, then to be RS, in order to easily compare the three levels. P values < 0.05 were considered significant.

## Results

We investigated the capability of two ICMS stimulation conditions to affect spike rate and temporal patterns of firing in a distant but interconnected cortical region. In healthy, ketamine-anesthetized rats, multi-site microelectrode probes were used to determine the effects of ICMS delivered to somatosensory cortical areas (FL and BF) on spontaneous spike firing rates, regularity of firing patterns, and stimulus-evoked spike firing in RFA.

### ICMS results in increased firing rate in a distant cortical area

First, mean firing rate (MFR) was compared between the initial basal period (Basal0) and subsequent basal periods (Basal1, Basal2, and Basal3) following each one-hour period of RS or ADS ICMS. Fig. 2A illustrates spike firing rates in a representative animal (ICMS in BF, ADS condition) demonstrating an overall increase of firing from Basal0 (top, left) to Basal3 (top, right). The increase in MFR occurred in each of the experimental conditions (ADSBF, ADSFL, RSBF, RSFL) but not in control experiments (CTRL) (Fig. 2B).

**Fig. 2.**
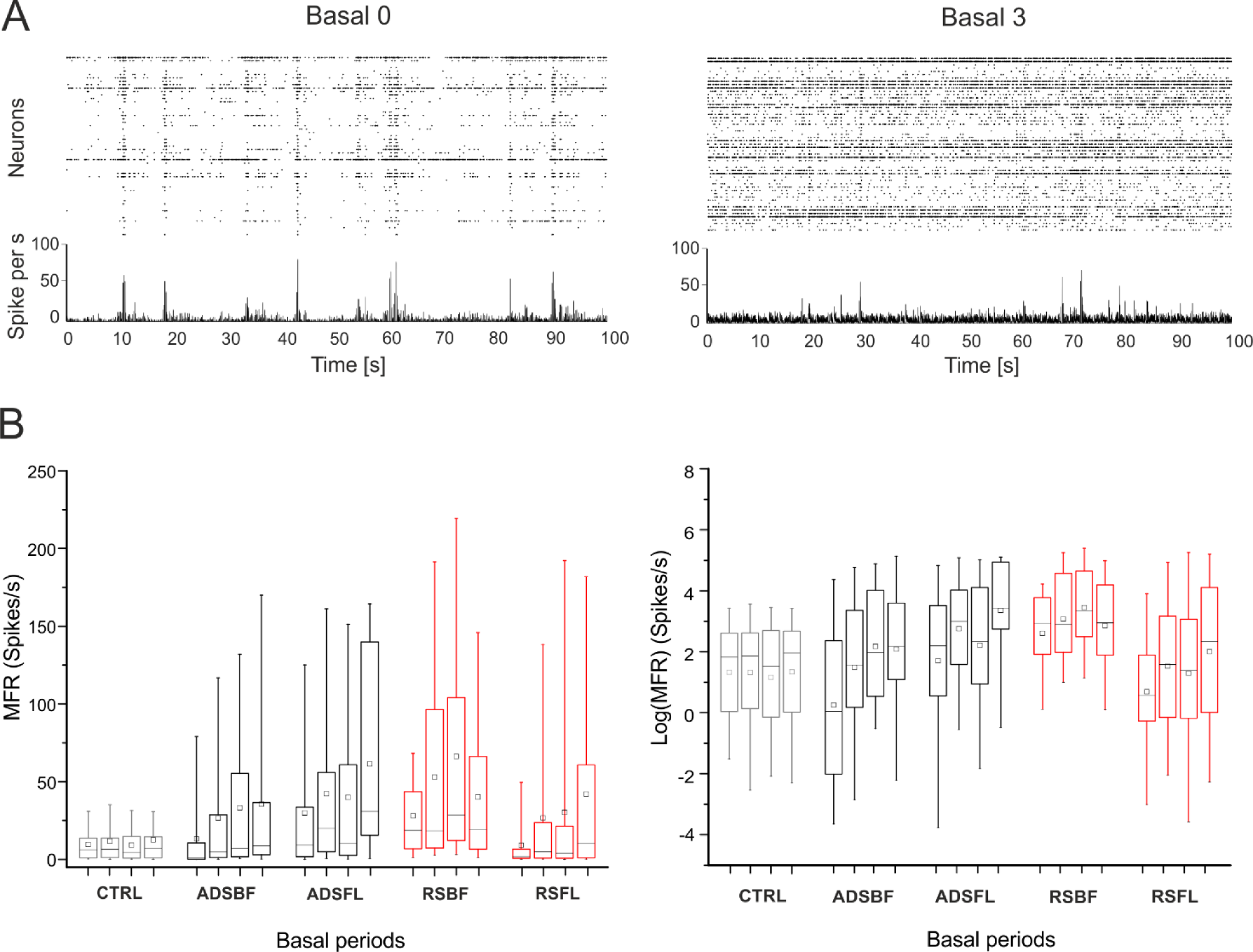
Firing rate analysis. (A) Spike rasters (dotted graphs, one row per neuron) and corresponding array-wide firing rate (line graphs) measured by summing all spikes detected on the entire array in 1-ms windows during 100 ms time frame of Basal0 (left) and Basal3 (right) for a single representative animal which underwent the ADS protocol in BF. (B) *Left* Quantitative representation of the Mean Firing Rate (MFR) distributions for each experimental group (CTRL, light grey; ADSBF, black; ADSFL, black; RSBF, red; RSFL, red) in each Basal phase (1 to 4). *Right* Representations of the Log(MFR) distributions of each experimental group and for each Basal period. Data are summarized in box plots, where the horizontal lines denote the 25^th^, the median and the 75^th^ percentile values and the whiskers denote the 5^th^ and the 95^th^ percentile values; the square inside the box indicates the mean of each dataset. Statistical analysis is reported in Table 2 and 3.

Table 2 contains the global results of hypothesis tests for the considered fixed effects. Main effects and third-order interactions of the effects are described in Table 3.

**Table 2:**
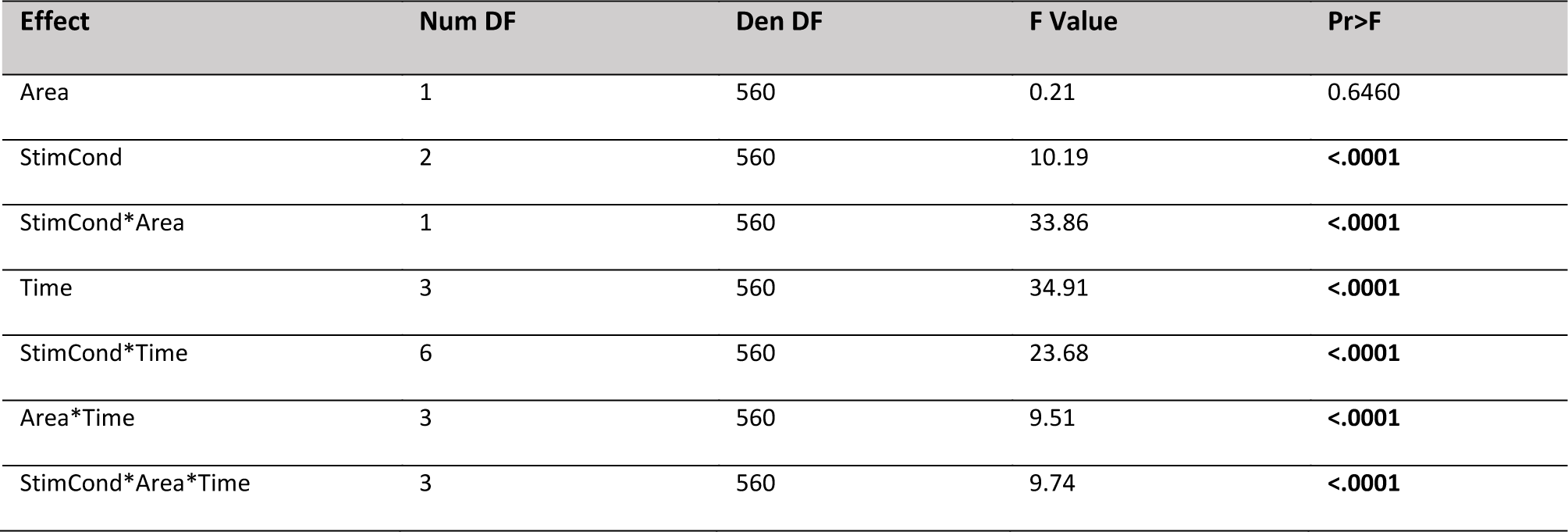
Results of general linear mixed effects model on the logarithmic firing rate log(MFR): hypothesis tests for the significance of each of the fixed effects considered. Type 3 Tests of Fixed Effects

| Effect | Num DF | Den DF | F Value | Pr>F |
| --- | --- | --- | --- | --- |
| Area | 1 | 560 | 0.21 | 0.6460 |
| StimCond | 2 | 560 | 10.19 | <.0001 |
| StimCond*Area | 1 | 560 | 33.86 | <.0001 |
| Time | 3 | 560 | 34.91 | <.0001 |
| StimCond*Time | 6 | 560 | 23.68 | <.0001 |
| Area*Time | 3 | 560 | 9.51 | <.0001 |
| StimCond*Area*Time | 3 | 560 | 9.74 | <.0001 |

**Table 3:**
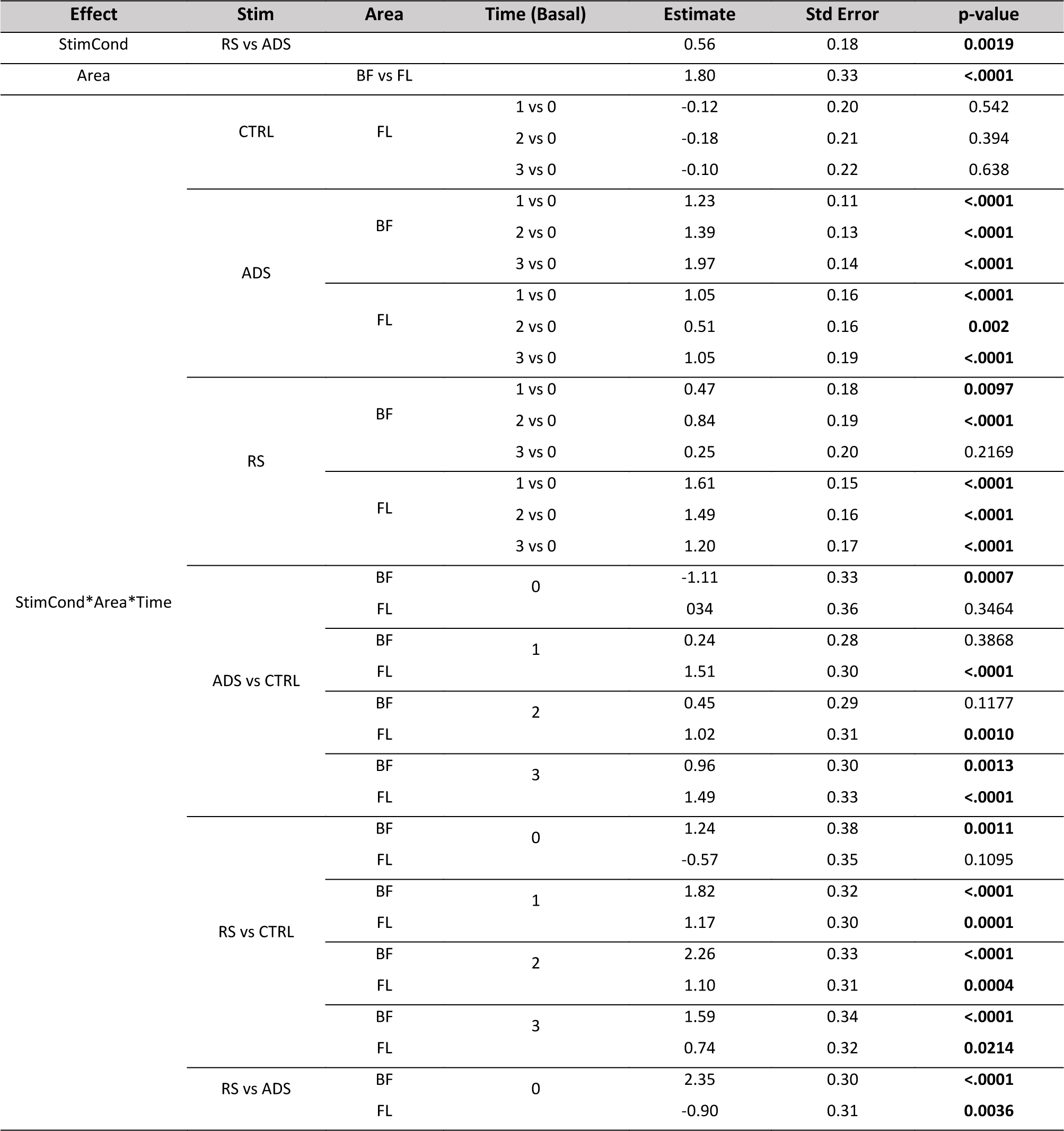
Results of the general linear mixed effects model on the logarithmic firing rate log(MFR): differences of Least Squares Means (the marginal means are estimated over a balanced population). Significant p-values are highlighted in bold. Note that for the control experiments (CTRL), we implanted a stimulating electrode in FL but, no stimulation was delivered.

Due to firing rate variability typically observed among different animals, as well as temporal fluctuations in firing rates in anaesthetized preparations, it was important to examine multiple post-stimulation basal periods in each animal. This approach provided not only greater statistical power, but allowed us to examine cumulative changes in firing rate in successive basal periods. In control (CTRL) experiments, no significant differences were found in MFR between any of the basal periods (Fig. 2B and Table 3). In contrast, MFR was significantly increased in 5 of 6 RS, and 6 of 6 ADS basal period comparisons. The only exception was the Basal3 vs. Basal0 comparison for RS in BF.

It is important to note that both ADS and RS generally induced a significant increase in firing rate in subsequent basal periods with respect to control experiments, regardless of any observed differences. As reported in Table 3 (see Table 3, *StimCond*Area*Time, ADS/RS vs CTRL)*, in the ADSBF group, MFR was significantly lower than in the CTRL group during Basal 0, but showed significantly higher MFR compared with CTRL in Basal3. For the ADSFL group, there were no significant differences with the CTRL group in Basal 0 but a significantly higher MFR in each of the subsequent basal phases. For the RSBF group, MFR was higher than the CTRL group for each the basal periods, while RSFL demonstrated a comparable MFR with respect to CTRL in Basal 0, but showed significant greater values for each of the subsequent phases. When considering the Effect *Area*, the stimulation of BF was more effective in increasing firing rate than the FL (see Table 3, *Area, BF vs FL*).

A table containing all the combinations of the third-order interaction *StimCond*Area*Time* is reported in the Supplementary Material section (see Table S1).

We also calculated the proportion of units (i.e., neurons) whose firing rates significantly increased, decreased or remained constant after the stimulation protocol (i.e., Basal1, Basal2, Basal3) with respect to the initial basal period of recordings (i.e., Basal0, Fig. 3A, 3B). Both stimulation conditions induced a significantly greater proportion of units that showed increased firing rates across basal periods (Fig. 3C) compared to the control group. No differences were observed in the proportion of units showing a decrease in MFR in the stimulation (Fig. 3D) vs the control experiments.

**Fig. 3.**
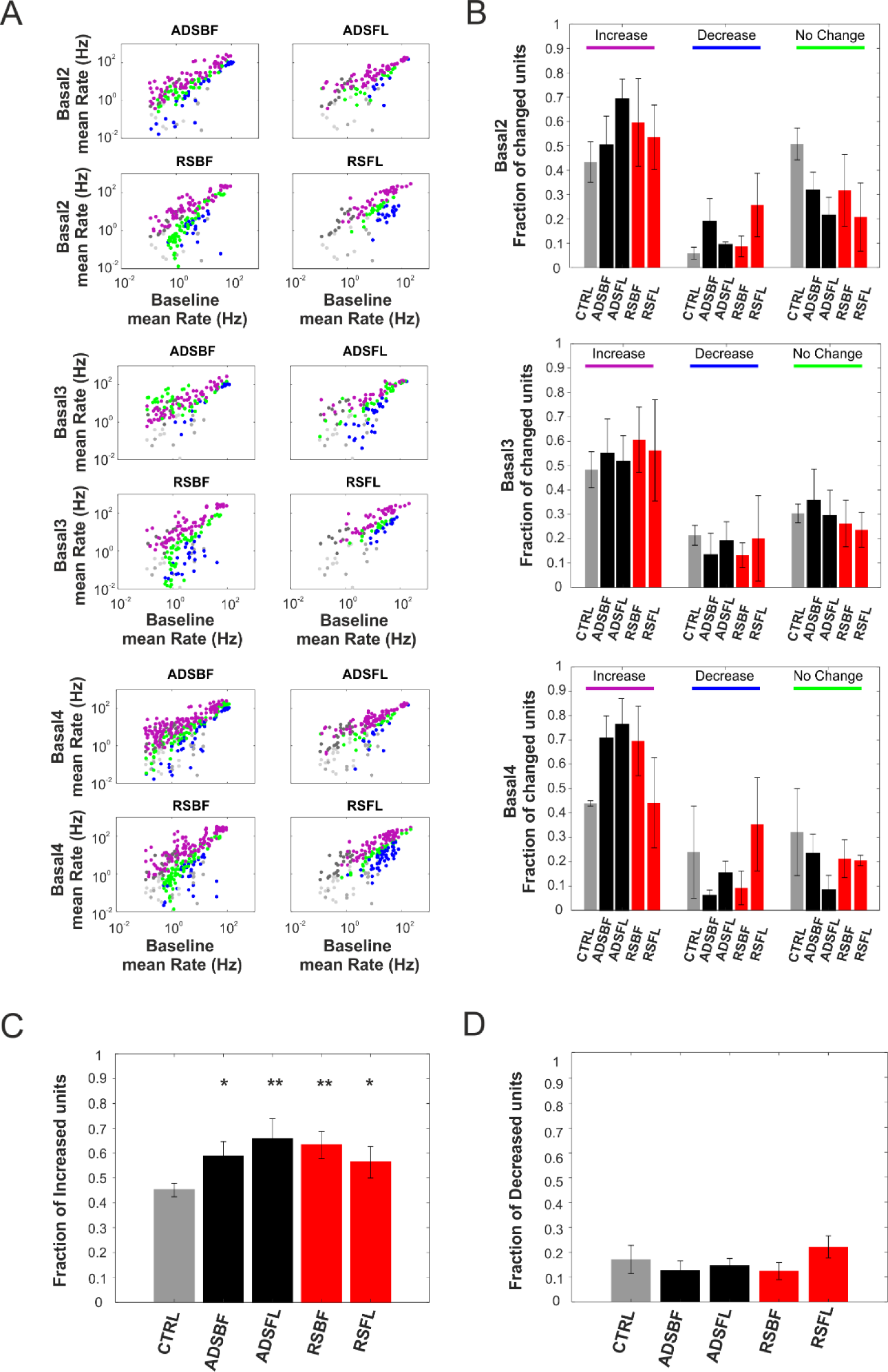
(A) Per unit correlation between baseline firing rates (x-axis, Basal0) and firing rates after each stimulation session (y-axis, Basal1, Basal2, Basal3) calculated for each group (ADSBF, ADSFL, RSBF, RSFL). Colors represent units that significantly increased (magenta), decreased (blue) or remained stable (green). Grey dots represent the correlation for the control group (CTRL). (B) Average fraction of units that significantly changed their firing with respect to the baseline period of recording (Basal0) calculated for all the five experimental groups (i.e. CTRL-light grey; ADSBF & ADSBF black; RSBF & RSFL red). (C) Average fraction of neurons which increased their firing rate across the five experimental groups. (D) Total fraction of neurons which decreased their firing rate. (*p < 0.05, relative to CTRL; one-way ANOVA with Dunnett’s multiple comparison test. Error bars represent SEM). Data are reported as mean ± SEM (standard error of the mean).

Overall these results demonstrate that, notwithstanding possible initial differences in MFR among groups, both RS and ADS stimulation conditions significantly increased the MFR of neurons recorded from RFA in the post-ICMS basal periods with respect to the initial basal period. ADS consistently showed an increase in MFR over time, in contrast to RS which showed an increase in MFR, but not for all of the experimental conditions and in sharp contrast to CTRL which never exhibited an increase in MFR. Increased MFR was accompanied by an increase in the proportion of units with increased firing rates in both RS and ADS stimulation conditions.

### ICMS modulates neuronal firing patterns in a distant cortical area

We used the LvR coefficient (Fig. 4A, cf. Materials and Methods), a metric of local variation of the inter-spike interval (ISI), to ‘classify’ the type of spike-firing pattern (e.g., ‘*Bursty’, ‘Random’* and ‘*Regular’)* of the recorded neurons in RFA during spontaneous activity. The purpose of this analysis was to understand whether the different ICMS stimulation conditions in the two areas affected spike-firing patterns. LvR distribution of neurons exhibited stable baseline firing patterns, with LvR values between ‘*Random’* and ‘*Bursty’* states (see Fig. 4B, ADSBF in Basal0, dotted line: LvR = 1.27±0.24, mean±SD).

**Fig. 4.**
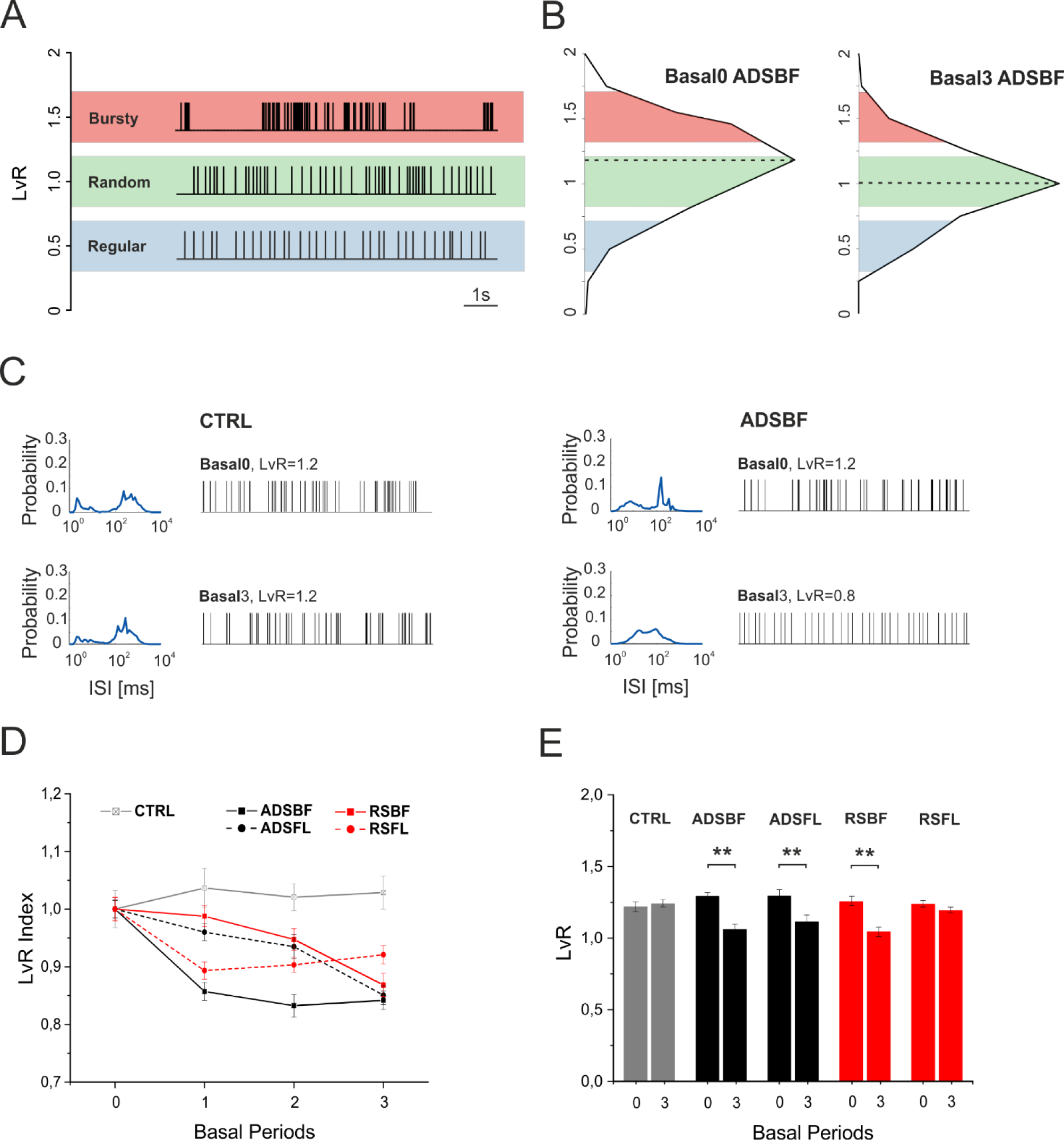
(A) Example spike sequences of representative neurons with LvR of 0.5 (Regular, red), 1 (Random, green) and 1.5 (Bursty, blue) taken from dataset ADSBF. (B) LvR distributions shown as histograms with a common bin size 0.25 and determined across all the subjects belonging from ADSBF during the first (Basal0, left) and last (Basal3, right) period of quiescence. Black dotted lines represent the median of the LvR distributions. (C) Distributions of a representative neuron’s interspike interval (ISI) and its sample firing pattern consisting of 100 consecutive ISIs belonging to the CTRL group (Left) and ADSBF group (right) respectively during Basal0 (top) and Basal3 (bottom). (D) Mean ± SEM trend of normalized LvR for each experimental condition. Each subject’s LvR value was normalized over the mean LvR value calculated during Basal0: statistical analysis is reported in Table 4 and 5. (E) Mean ± SEM LvR comparison between Basal0 and Basal3 for all the experimental conditions (CTRL, grey; ADSBF and ADSFL, black; RSBF and RSFL, red). **p < 0.01; unpaired, two-tailed Student’s t-test.

Table 4 contains results of hypothesis tests for each of the considered fixed effects. It shows that the effect *StimCond* and the combinations *StimCond*Area* and *StimCond*Area*Time* were not significant. The global effect *Area* reached significance (p=0.0497), while *Time* and the other interactions were clearly significant.

**Table 4:**
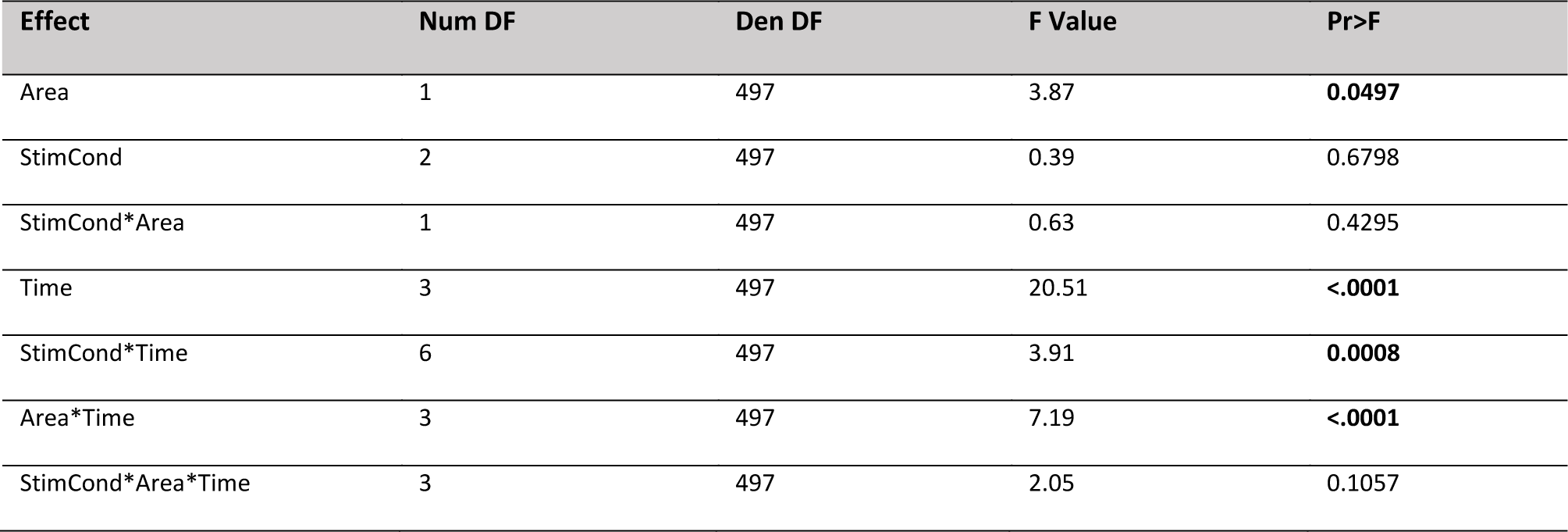
Results of general linear mixed effects model on LvR: hypothesis tests for the significance of each of the fixed effects considered. Type 3 Tests of Fixed Effects.

**Table 5:**
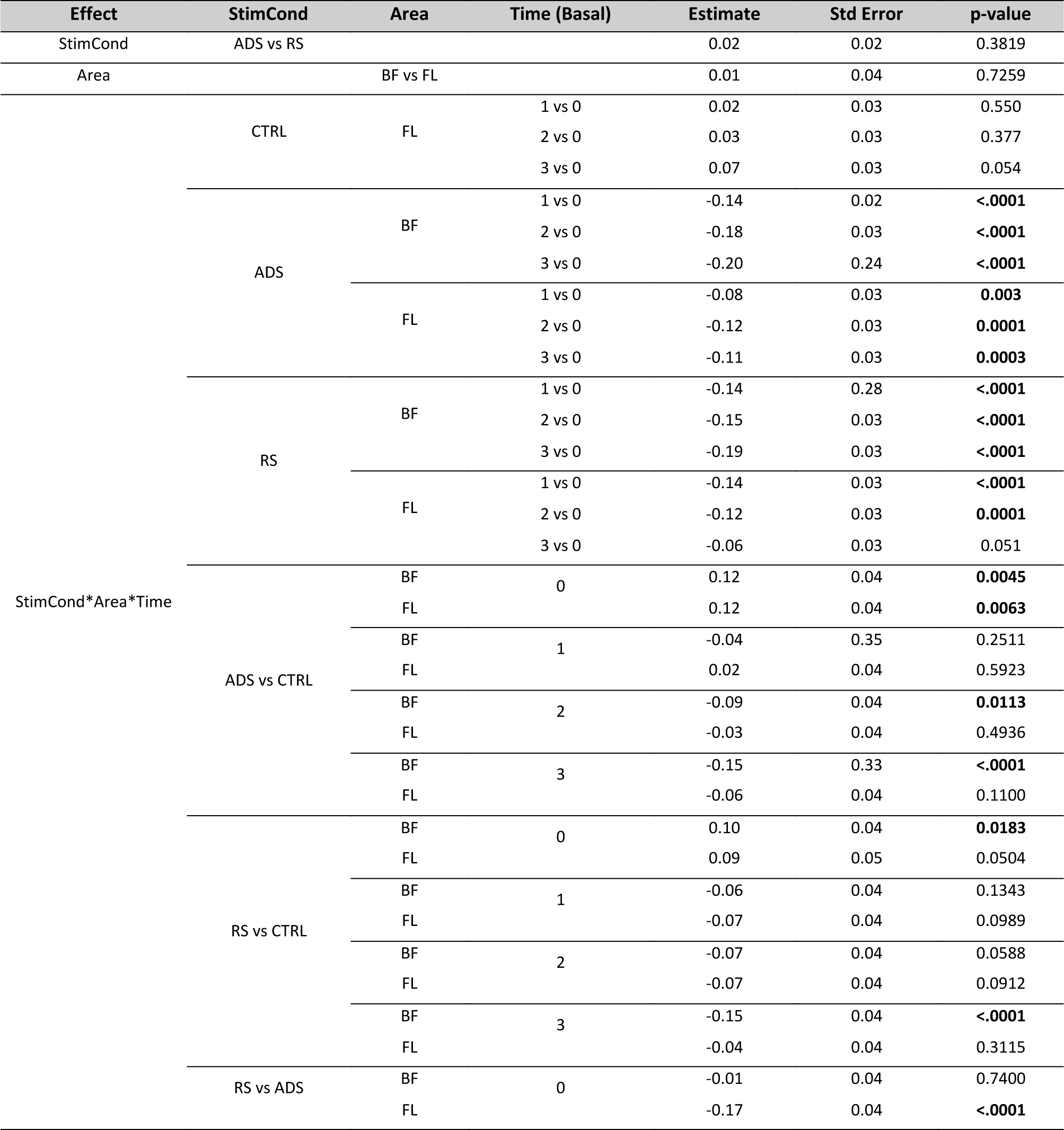
Results of general linear mixed effects model on LvR: differences of Least Squares Means (the marginal means are estimated over a balanced population). Significant p-values are highlighted in bold.

No statistical difference was found for LvR values comparing the two stimulation conditions (see Table 5, Effect *StimCond*, ADS vs RS). The contrast between FL and BF indicates that there were no significant differences between the two stimulated somatosensory areas (Table 5, Effect *Area*, BF vs FL). Table 5 also reports how both ADS and RS altered the firing patterns across time. As expected, LvR was not affected in the CTRL experiments (Fig. 4C-E and Table 5, *StimCond*Area*Time* CTRL). The main result is that ICMS, either RS or ADS, generally induced a strong *decrease* in LvR, moving activity from the ‘*Bursty* condition during Basal 0 towards the ‘*Random’* state of firing in the last basal period (Fig. 4B, ADSBF in Basal3, dotted line: LvR = 1.07±0.02, mean±SD and Fig. 4D-E, Table 5). This effect was observed in 5 of 6 basal period comparisons using RS and 6 of 6 basal period comparisons using ADS. The one exception was that no change in LvR was observed between Basal3 and Basal0 in the RSFL group (Table 5).

As was observed for MFR, there were differences between groups in LvR even during the initial basal period (Basal0). However, the significant changes that occurred in LvR in the post-ICMS basal periods invariably were decreases. For example, ADSBF LvR was significantly higher than CTRL during Basal 0, but this difference disappeared in Basal1. ADSBF showed a significantly lower LvR than the CTRL in Basal2 and Basal3, indicating an overall decrease of LvR for ADSBF group with respect to CTRL. LvR in the ADSFL group was significantly higher than the CTRL in Basal0, but the difference disappeared in all subsequent basal phases. The RSBF group was higher than the CTRL in Basal0, but the difference disappeared in Basal1 and Basal2. Finally, RSBF was significantly lower than CTRL during Basal3. RSFL was found to be not statistically different from CTRL in any of the basal periods. A table containing all of the combinations of the effect *StimCond*Area*Time* is reported in the Supplementary Material section (see Table S2).

### RS and ADS exhibit different effects on stimulus-associated action potentials in a distant cortical area

Evoked action potentials (in RFA) in response to ICMS were analyzed by discriminating the spiking activity in the 28ms after each S1FL or S1BF stimulus pulse (Fig. 5C) using post-stimulus time histograms (PSTH, cf. Materials and Methods). Due to differences in the dynamics of the evoked electrical artifacts in different animals, we used an adaptive-length blanking window (from 4 to 6ms, Fig. 5A). As shown in Fig. 5B, the interstimulus intervals (ISIs) used for the RS groups comprised a range of values comparable to those of the ADS groups. Interestingly, the ISI distributions for ADS was stable over repeated stimulation trials (see Fig 5B, left). There was, however, a bias toward ~200ms interstimulus intervals in the ADS group, whereas the RS intervals exponentially decreased across the range (Fig. 5B).

**Fig. 5.**
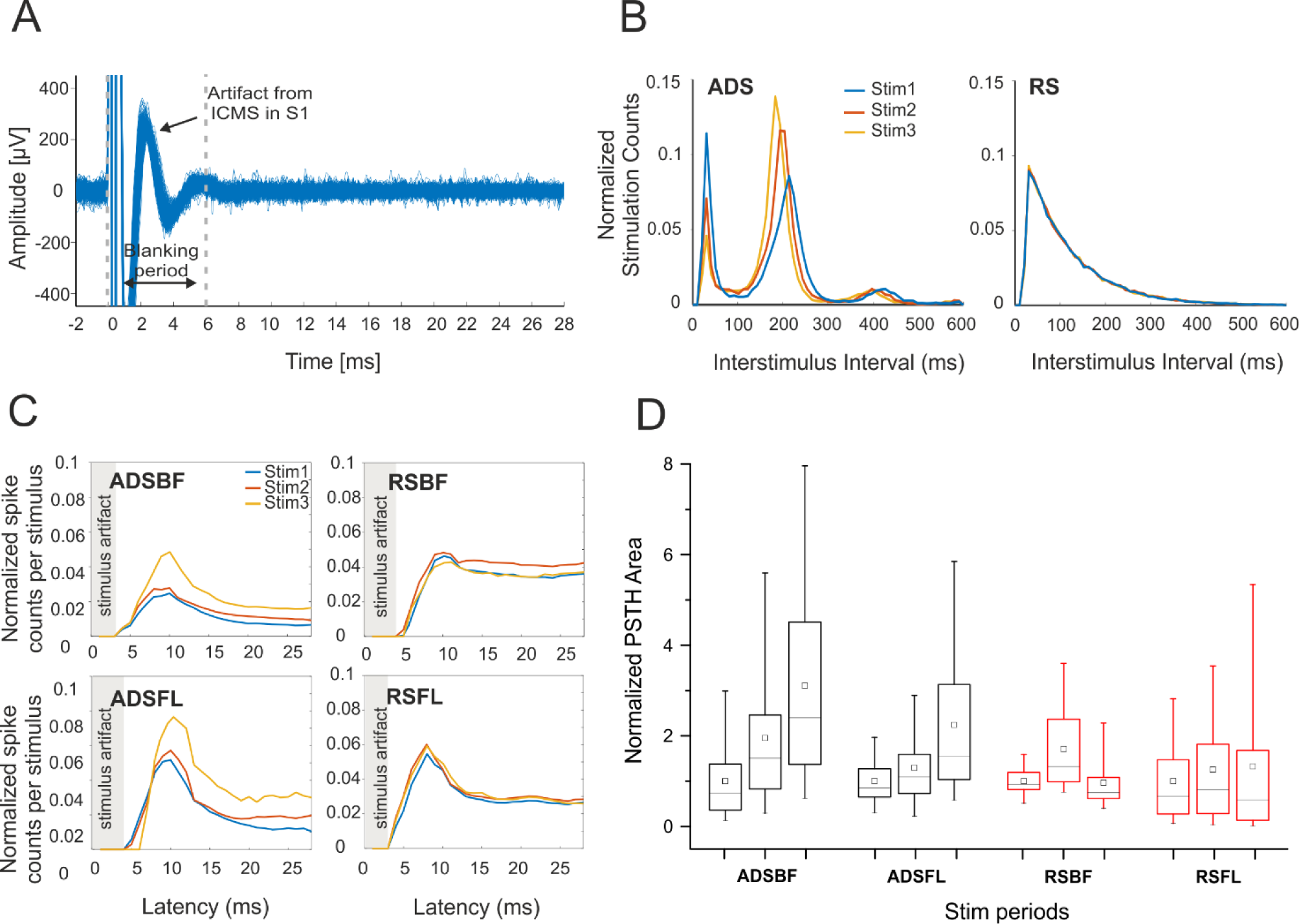
(A) Sample trace of recordings from RFA showing stimulus artifacts from ICMS delivered to S1BF. Blanking period used for the analysis is delimited by the grey dotted lines. A total of 100 superimposed traces are shown. (B) Stimulation interval distribution (Interstimulus intervals, 600-ms) in both one ADS and RS subject during the three stimulation sessions (Stim1, Stim2 and Stim3). Stimulation counts have been normalized to session length. (C) Post-stimulus spiking histograms derived from neural recordings in RFA on the three stimulation sessions for the four ICMS groups respectively (ADSBF, ADSFL, RSBF, RSFL). Histograms portray the average number of action potentials discriminated from the neural recordings within 1-ms bin. Data pooled across subjects for each group and normalized to the total number of either ADS or RS specific events. (D) Normalized PSTH areas for the four groups (ADSBF and ADSFL, black; RSBF and RSFL, red) of ICMS in the three stimulation phases (Stim1, Stim2, Stim3). Each subject’s PSTH area were normalized over the mean area calculated in Basal0 to show data trends over time. Statistical analysis is reported in Table 6 and 7.

Table 6 contains results of hypothesis tests for each of the considered fixed effects. It indicates that the effects *Area, Time* and their interactions induced significant changes in the evoked activity. However, the fixed effect *StimCond* was not significant (see Table 6, Effect *StimCond*).

**Table 6:** Results of general linear mixed effects model on PSTH: hypothesis tests for the significance of each of the fixed effects considered. Type 3 Tests of Fixed Effects.

| Effect | Num DF | Den DF | F Value | Pr>F |
| --- | --- | --- | --- | --- |
| Area | 1 | 299 | 16.71 | <.0001 |
| StimCond | 1 | 299 | 0.10 | 0.7469 |
| StimCond*Area | 1 | 299 | 45.42 | <.0001 |
| Time | 2 | 299 | 30.73 | <.0001 |
| StimCond*Time | 2 | 299 | 17.58 | 0.0008 |
| Area*Time | 2 | 299 | 16.55 | <.0001 |
| StimCond*Area*Time | 2 | 299 | 3.32 | 0.0375 |

**Table 7:**
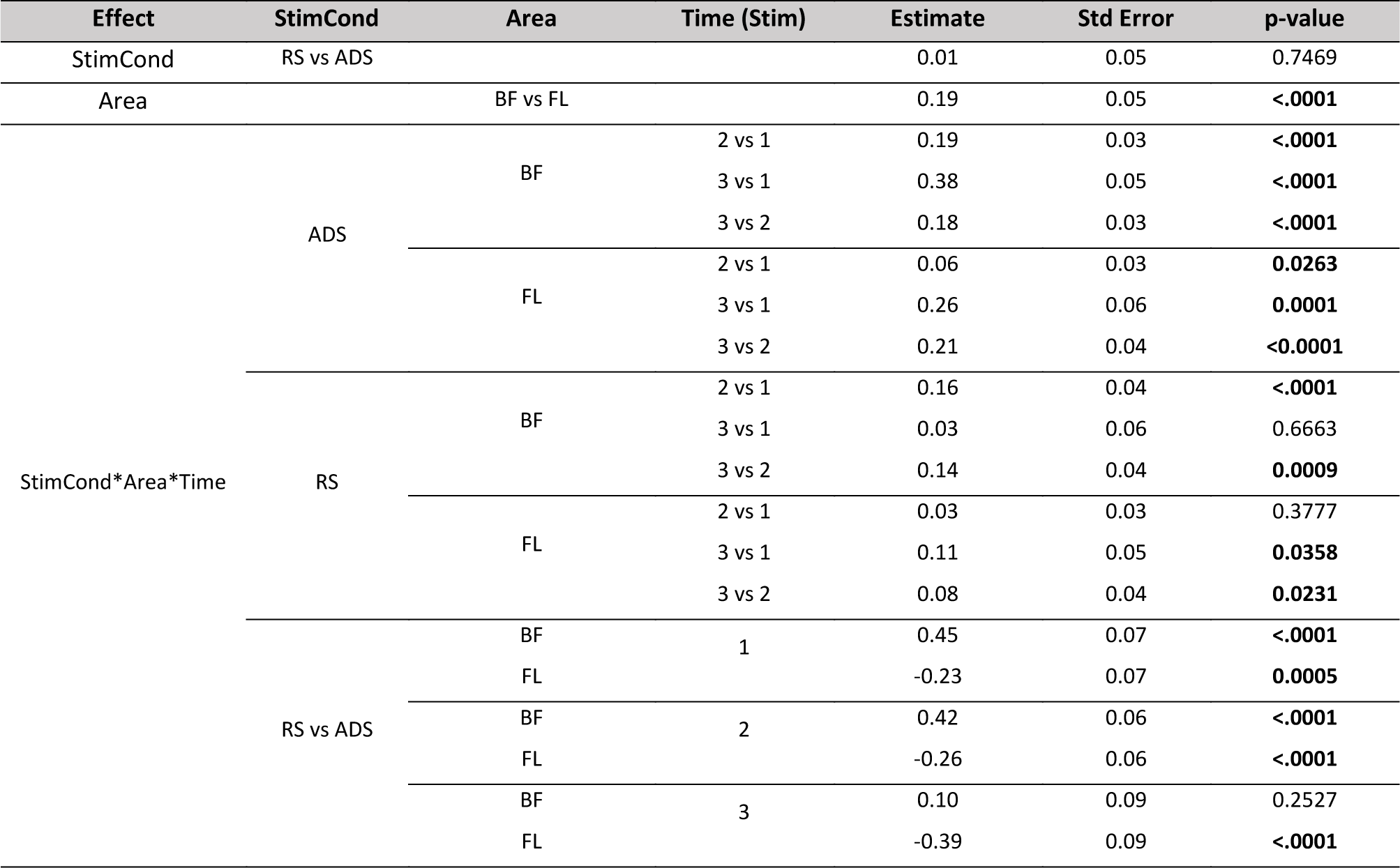
Results of general linear mixed effects model on PSTH: differences of Least Squares Means (the marginal means are estimated over a balanced population). Significant p-values are highlighted in bold.

Our analysis indicated that ICMS in BF was significantly more effective in evoking short-latency spikes (<28 ms) than in FL (Table 7, Effect *Area*, BF vs FL). Examining differences between the two stimulation conditions, it appeared that ADS had a more consistent, cumulative effect in eliciting evoked spikes over time (see Table 7, Effect *StimCond*Area*Time*, ADS, both BF and FL). This is evident in Fig. 5C and D, showing a progressive increase in short-latency spike counts and PSTH area, respectively in ADS. Changes in evoked spike activity with RS were smaller and inconsistent. Statistically significant differences between RS and ADS were found in 3 of 3 of the FL stim period comparisons and 2 of 3 BF stim period comparisons (Table 7).

Regarding the stimulus location, we observed that ADSFL consistently evoked more post-stimulus spikes than RSFL. A table containing all the combinations of the effect *StimCond*Area*Time* is reported in the Supplementary Material section (see Table S3).

## Discussion

For several decades, electrical microstimulation has been an important tool for investigating neural circuits, demonstrating evidence of cortical map plasticity, and for therapeutic neuromodulation [42–44]. Despite its widespread use, our understanding of its effects beyond local depolarization of neuronal membranes remains limited. While the behavioral outcomes of microstimulation in sensory and motor regions of the brain have been characterized extensively [45, 46], few studies have examined the long-term effects of repetitive microstimulation on neuronal activity in the broader network of interconnected brain regions [47]. Those that have examined distant effects of microstimulation have focused primarily on the alteration in neuronal activity in primary motor cortex (M1) induced by the stimulation of the subthalamic nucleus (STN) as a model to understand the effects of DBS therapy in Parkinson’s disease [44, 48–51]. As adaptive, or closed-loop, microstimulation modalities are increasingly being explored as potential options for therapeutic applications [28, 52], it is important to understand how various stimulation patterns differentially alter both spontaneous and stimulus-evoked neuronal activity in interconnected regions of the brain. The specific aim of the present work was to characterize the neurophysiological effects of random and activity-dependent ICMS on healthy cortical networks. Indeed, our study indicates ICMS is able to alter, within hours, the firing characteristics within a distant, but connected, cortical area (RFA).

To determine the impact of focal electrical microstimulation on distant cortical regions in healthy anesthetized rats within single recording sessions, we applied either randomized ICMS (i.e., open-loop) or activity-triggered ICMS (i.e., closed-loop) to one of two somatosensory cortical areas (forelimb or barrel field) while recording resultant neuronal activity in the RFA, a premotor cortical area. These regions were chosen due to their known intracortical connections and our ability to alter synaptic efficacy in the target pathways in a previous study [28]. We found that ICMS in somatosensory areas induced an increase in spontaneous firing rates in RFA when compared to non-stimulated controls (Fig. 2 and Fig. 3, Table 3). Further, we observed a reduction in the LvR values, indicating a shift towards more random interspike intervals of recorded units within RFA (Fig. 4E, Table 5). Finally, we found an increase in the stimulus-evoked activity in RFA (Fig. 5, Table 6, 7). While both forms of stimulation induced increases in spontaneous firing rates and decreases in LvR, effects were marginally more consistent with ADS. A more pronounced difference between the two stimulation conditions was found in the ability to evoke short-latency spikes (<=28 ms) in RFA. Increases in evoked spikes as a result of ADS were progressive over multiple stimulation periods, and significantly different from results of RS.

Both RS and ADS modulated spontaneous firing rates and patterns within RFA in a similar manner. Both resulted in decreased LvR which was initially associated with a mixed ‘*Random’-‘Bursty’* intrinsic firing pattern, indicating a shift towards a ‘*Random’* state of firing at the end of the treatment (see Figure 4E). Regarding the role of the stimulus location, we found that stimulation from BF was more effective than FL in increasing the spontaneous firing rate in RFA (see Table 3, effect *Area*, BF vs FL). Examining the evoked response, stimulation from BF was also more effective in directly evoking action potentials in RFA (see Fig. 5 C, D and Table 6, effect *Area*, BF vs FL).

Given the reciprocal cortico-cortical connections between RFA and the two somatosensory areas, these results were unexpected [53]. Other cortical (and subcortical) structures undoubtedly play a role in these distant effects of repetitive microstimulation (cf. Fig. 1). The primary motor cortex (caudal forelimb area or CFA in rats), has dense reciprocal connections with RFA as well as the somatosensory areas. However, if CFA activity was modulated in the present paradigm, one would expect that the influence would be greater with S1FL stimulation compared to S1BF stimulation, due to the important role in sensorimotor integration mediated by CFA-S1FL connections. This hypothesis will need to be verified by simultaneously measuring spike activity from CFA.

Both ICMS protocols were able to induce changes in firing rate with respect to non-stimulated animals (CTRL) where no changes were observed. Interestingly, even if there was an initial difference between ADS and RS groups during the baseline period of recording, ADS was invariably able to increase firing rates over time, compared with RS. No differences were found in the LvR analysis (Table 5, Effect *StimCond*, RS vs ADS) but ADS more reliably displayed lowered LvR over time (c.f. Table 5, Effect *StimCond*Area*Time*, Time 4 vs 1 for all the groups).

More interestingly, ADS, and not RS, facilitated progressive increases of stimulus-associated activity over time (see Effect *StimCond*Area*Time* for ADS and RS in Table 7) suggesting that the pairing of neural activity and stimulation may lead to stronger associations over prolonged stimulation sessions. This may result from a number of factors. One potential mechanism that allows ADS to be more effective than RS is that ADS is thought to utilize a Hebbian-based spike-timing method similar to the nervous system’s natural mechanism to promote learning and memory which is typically effective in inducing long term plasticity [24, 28].

While these studies provide important evidence for the effects of electrical microstimulation on the broader neuronal network, it is important to consider the limitations imposed by the ketamine-anesthetized preparation. An anesthetized preparation has numerous advantages for the present investigation, since the state of the animal and the associated neurophysiological set-up is relatively stable over several hours. While it is technically feasible to conduct these studies in awake, ambulatory animals, substantial variability in spike activity is introduced by the sensorimotor activities of the animals. However, ketamine is widely known as a noncompetitive N-methyl d-aspartate receptor antagonist, and can modulate other receptors or channels such as the GABAa receptor. As a result, ketamine has diverse and temporally complex effects on neuronal activity [54–56]. For example, under ketamine anesthesia, different cell types in the hippocampus show differential effects in firing rate and synchrony [57]. Ketamine causes enhanced gamma oscillations acutely, but decreased network gamma oscillations with chronic [58] administration. Thus, while ketamine anesthesia undoubtedly had some effect on neuronal firing in the present study, the changes in MFR, LvR and evoked spikes are thought to be largely independent of the anesthetic state. This hypothesis will need to be verified in awake, ambulatory animals.

In summary, RS and ADS protocols both induce changes in the recorded activity in RFA. It is clear that focal electrical stimulation has the ability to alter activity in remote brain regions not directly influenced by the current spread from the electrode. The closed-loop condition (ADS) can effectively be considered as more reliable in its ability to alter evoked responses and thus in modulating cortico-cortical connectivity in the rat brain within a single recording period. Closed-loop stimulation for therapeutic applications in the human brain is still uncommon. However, many similar approaches are already being tested for epilepsy, in Parkinson disease and in animal models of spinal cord injury [59–62]. Other potential clinical applications based on closed-loop ICMS treatments include stroke, focal TBI, and surgical resections. Although the beneficial effect of these approaches in humans is still not clear, we propose that ADS could be used to modulate cortical state and connectivity by steering neuroplasticity after injury. Additional studies need to be performed to determine the precise parameters and characteristics related to these alterations.

## Acknowledgments

Project funded by the Italian Ministry of Foreign Affairs and International Collaboration (MAECI), Directorate General for Country Promotion, as a high-relevance bilateral project within the Italy-USA. Partial support was also provided by NIH R01NS030853 (RJN) and NIH R03HD094608 (DG). The authors thank Giacomo Siri from the University of Genova for his support in the statistical model development and Caleb Dunham from the University of Kansas Medical Center for technical assistance with spike analysis and data management.

## Author contributions

AA, RJN, MC and DG designed the study. AA, GVD, MM and DG performed the experiments. AA, VP, MM, and MC designed the pipeline for the data analysis. AA performed the analysis. AA, MPR, VP performed the statistical analysis. AA, MC, DG and RJN wrote the paper. All the authors have read and approved the manuscript.

## Supplementary Materials

**Table S1:**
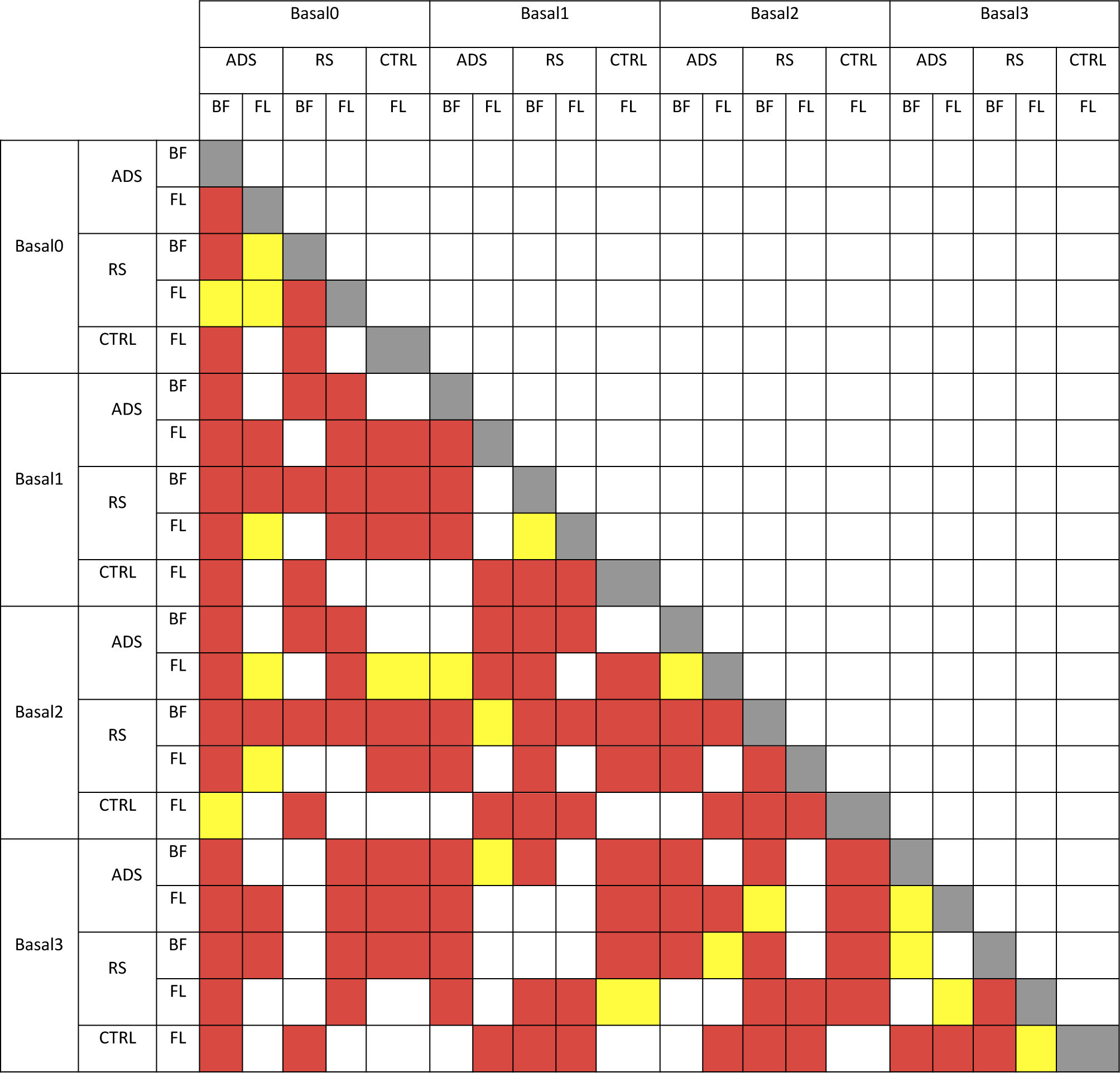
Results of general linear mixed effects model on the logarithmic firing rate log(MFR): differences of Least Squares Means (the marginal means are estimated over a balanced population) calculated for the *Stim*Area*Time* effect. Significant p-values are colored: p-value .01-.05 (weak evidence) yellow, p-value <.01 (very strong evidence) red.

**Table S2:**
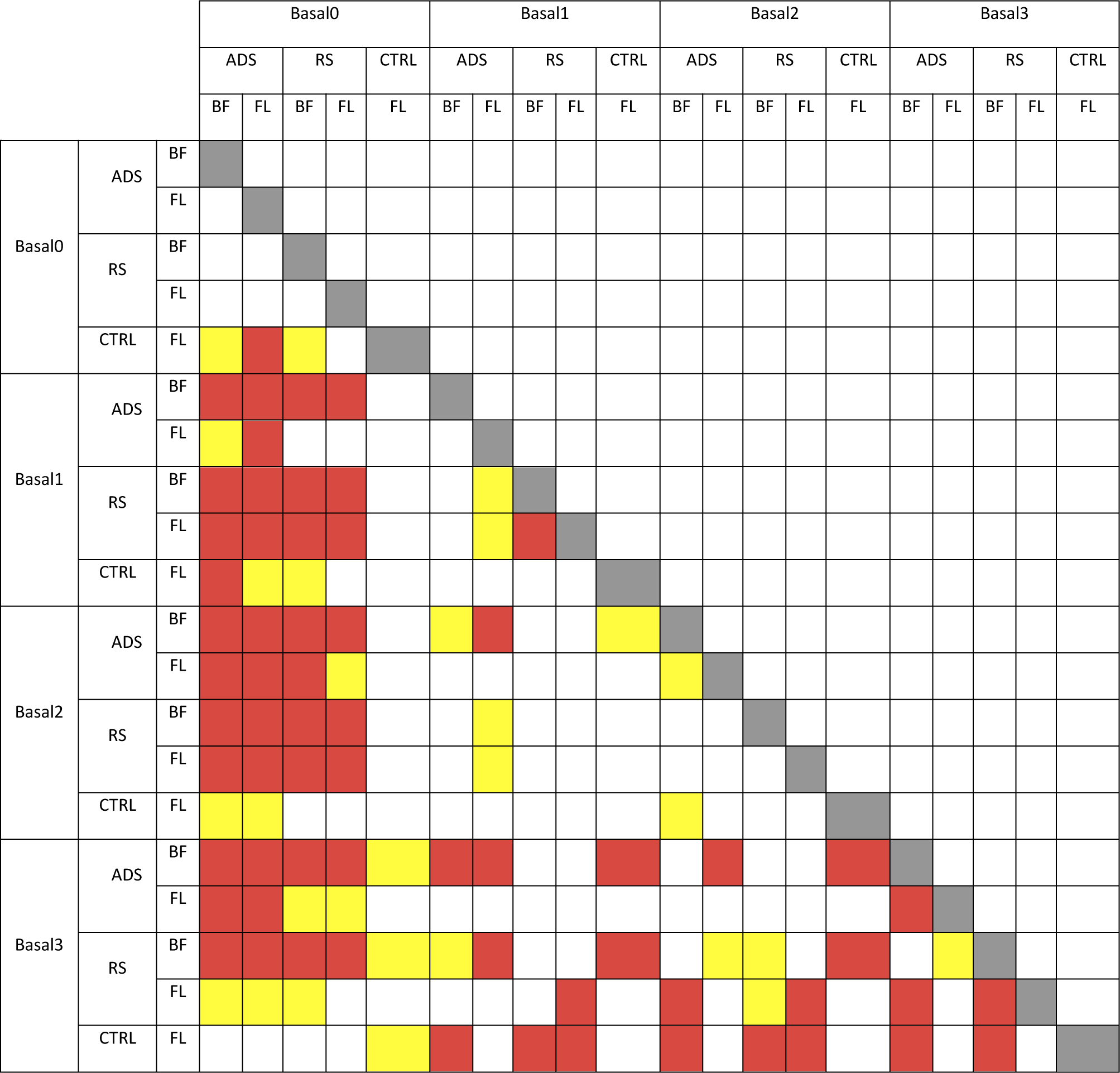
Results of the general linear mixed effects model on the LvR: differences of Least Squares Means (the marginal means are estimated over a balanced population) calculated for the *Stim*Area*Time* effect. Significant p-values are colored: p-value .01-.05 (weak evidence) yellow, p-value <.01 (very strong evidence) red.

**Table S3:**
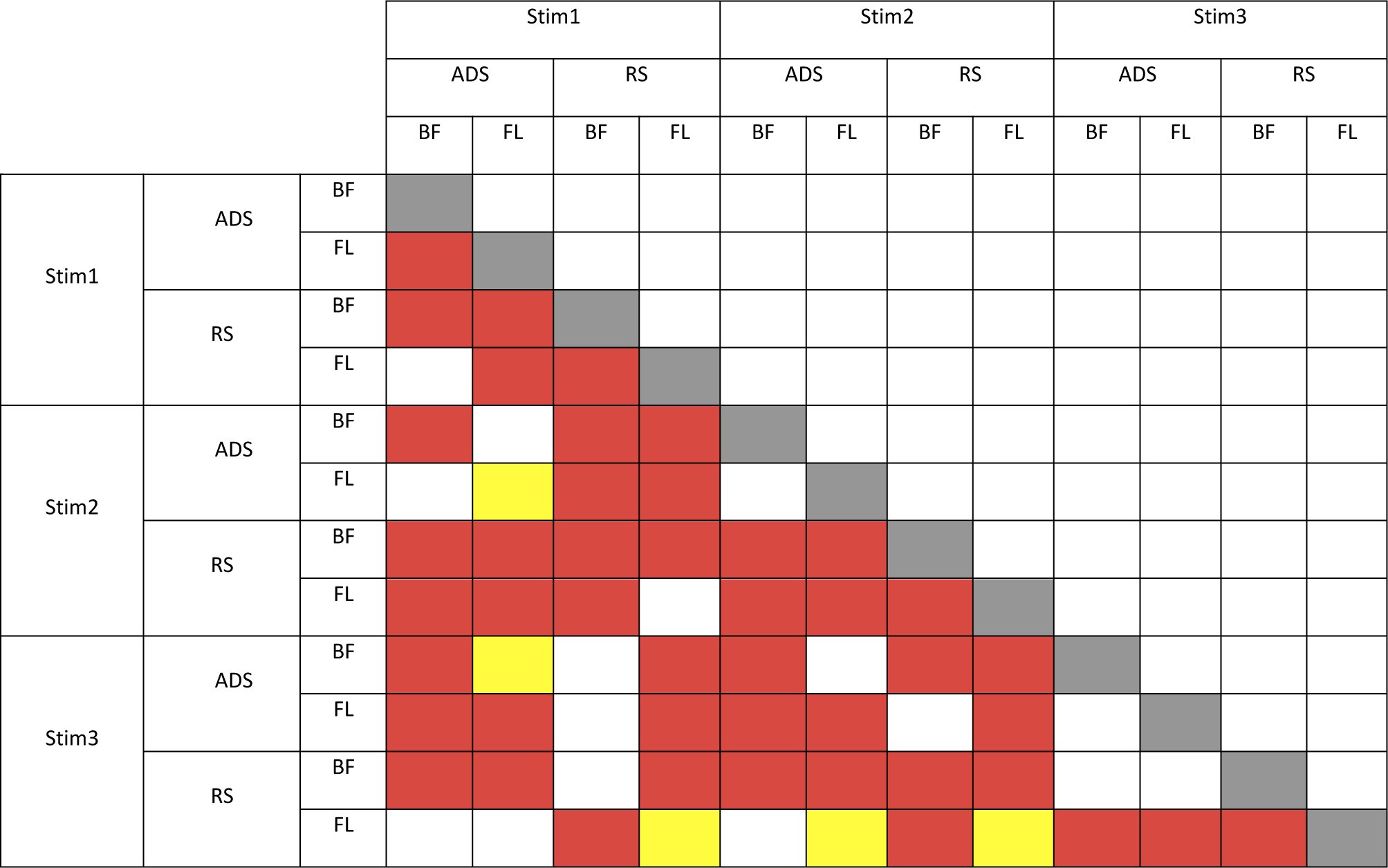
Results of the general linear mixed effects model on the PSTH: differences of Least Squares Means (the marginal means are estimated over a balanced population) calculated for the *Stim*Area*Time* effect. Significant p-values are colored: p-value .01-.05 (weak evidence) yellow, p-value <.01 (very strong evidence) red.

## Abbreviations

ADS: Activity Dependent Stimulation
BF: barrel field
CTRL: control
FL: forelimb area
ICMS: intracortical microstimulation
LvR: Local variation compensate for Refractoriness
MFR: mean firing rate
REML: restricted maximum likelihood
RFA: rostral forelimb area
RS: Random Stimulation
tDCS: transcranial direct current stimulation
TMS: transcranial magnetic stimulation

